# DECODING SHAPE ICONICITY OF WORDS AND PSEUDOWORDS: AN fMRI STUDY

**DOI:** 10.64898/2026.05.15.725463

**Authors:** G. Vinodh Kumar, Simon Lacey, Lynne C. Nygaard, K. Sathian

## Abstract

Iconicity refers to systematic links between word form and meaning. Although evidence for iconicity in natural language continues to grow, its neural basis remains unclear. Using functional magnetic resonance imaging (fMRI) and multivariate pattern analysis (MVPA), we examined iconic shape associations of auditory real words and pseudowords. The pseudowords were matched to the real words in phonemic and phonotactic properties, differing primarily in the absence of learned semantic representations. Participants listened to each item and judged whether it sounded rounded or pointed. Searchlight MVPA revealed significant decoding for both stimulus types. For real words, significant decoding was observed in regions associated with visual imagery (bilateral precuneus), in addition to regions associated with spectral processing (right superior temporal gyrus and sulcus extending into the parietal operculum). For pseudowords, significant decoding was observed primarily in regions associated with acoustic–phonetic processing (bilateral superior temporal gyri). Multivariate classifiers trained to distinguish rounded from pointed real words also distinguished pseudowords, and vice versa, in bilateral superior temporal regions, indicating shared information distinguishing rounded and pointed items across lexical conditions. Together, these findings suggest that similar acoustic–phonetic processing contributes to iconic shape associations for both real words and pseudowords. The additional involvement of imagery-related regions for real words suggests that established word meanings may also support access to perceptual information during shape judgments.

## 1. INTRODUCTION

Language is increasingly recognized to exhibit systematic, non-arbitrary alignments between word form and word meaning, a phenomenon known as iconicity. For example, onamatopoeic words such as *buzz* and *thud* resemble the sounds they denote, while pseudowords like *kiki* and *bouba* evoke non-auditory sensory properties, such as pointed and rounded shapes, respectively (Ramachandran & Hubbard, 2001). While natural language has traditionally been characterized as relying predominantly on arbitrary sound-meaning associations (Joseph, 2015; de Saussure, 1916), accumulating evidence indicates that arbitrariness and iconicity coexist as complementary properties of language, rather than being mutually exclusive (Campbell et al., 2025; Lockwood & Dingemanse, 2015; Perniss et al., 2010). However, the neural basis of iconicity and its role in sound-to-meaning mappings in natural language remain unclear.

Our understanding of the psychophysical and cognitive processes that mediate iconic sound-to-meaning mappings has been greatly advanced through the use of pseudowords, which allow investigation of iconicity while minimizing contributions from established semantic associations. Behavioral work has consistently identified properties associated with iconic mappings between sound and meaning via different analyses, including acoustic (Knoeferle et al., 2017; Kumar et al., 2025; Lacey et al., 2020, 2026), phonetic (Lacey et al., 2026; McCormick et al., 2015; Nuckolls, 1999; Sapir, 1929), and phonemic (Johansson et al., 2020; Lacey et al., 2026; Sidhu & Pexman, 2018). Each of these analyses reveal that specific spectro-temporal or other acoustic properties, phonetic features and phonemes systematically characterize associations between sound and meaning across different semantic domains such as shape and arousal (Kumar et al., 2025; Lacey et al., 2026), or social dominance (Auracher, 2017). Importantly, these effects are remarkably robust: they are observed across developmental stages (Maurer et al., 2006; Ozturk et al., 2013), diverse languages and geographical contexts ( Dingemanse et al., 2015; Blasi et al., 2016), and cultural and writing systems (Bremner et al., 2013; Ćwiek et al., 2022). This consistency suggests that iconicity reflects a fundamental aspect of language processing rather than being a language-specific or culturally constrained phenomenon.

In parallel with the behavioral literature, converging neuroimaging evidence indicates that iconicity is supported by distributed neural processes. Functional magnetic resonance imaging (fMRI) studies of onomatopoeic and Japanese mimetic words have implicated superior temporal regions in iconic processing (Hashimoto et al., 2006; Kanero et al., 2014). Extending this work to ordinary spoken nouns, congruence between the arousal conveyed by a word’s sound and its meaning has been associated with changes in responses of the left amygdala and its functional coupling with temporal and frontal regions (Aryani et al., 2019). Studies focusing specifically on sound-shape associations have used crossmodal paradigms, either implicit association (McCormick et al., 2022; Peiffer-Smadja & Cohen, 2019) or explicit congruency judgment tasks (Barany et al., 2023). Univariate analyses show that sound-shape congruency modulates blood oxygenation level-dependent (BOLD) responses across multiple regions: incongruent pairings tend to elicit greater activity in regions associated with phonological processing, attentional control, and crossmodal conflict (McCormick et al., 2022; Peiffer-Smadja & Cohen, 2019), whereas congruent pairings are associated with processing in auditory and higher-order visual regions (Barany et al., 2023). Multivariate pattern analyses (MVPA) found that iconic shape distinctions can be decoded in language-related regions and visual cortex (Barany et al., 2023). Taken together, the available neuroimaging evidence suggests that iconic mappings engage regions involved in language, attention, and multisensory processing.

Much of the neuroimaging work on sound-shape associations has relied on crossmodal paradigms involving congruent and incongruent pairings of auditory pseudowords and visual shapes, often using engineered pseudowords designed to elicit rounded-pointed distinctions. Although such stimuli are well suited for assessing the influence of phonological content on iconic mapping while minimizing confounds from semantic knowledge (Lacey et al., 2026; Kumar et al., 2025; Johansson et al., 2020; McCormick et al., 2015; Nuckolls, 1999), they pose a fundamental limitation. Particularly, engineered pseudowords do not capture the representational complexity of natural language where acoustic and phonological cues are embedded within established semantic representations and prior experience. Given the existing evidence for iconicity in real words (Campbell et al., 2025; Dingemanse et al., 2015; Perniss et al., 2010; Winter et al., 2023), it is important to determine the neural mechanisms that underlie iconicity in natural language. Specifically, it remains unclear whether the neural patterns observed with engineered pseudowords generalize to real words, or whether real-word responses are modulated solely by semantic knowledge. Addressing this gap is essential for understanding the neural mechanisms that underlie iconic associations during natural language comprehension.

The central question of this study is whether the neural mechanisms supporting iconic associations are shared between pseudowords and real words. If iconic associations are primarily grounded in acoustic or phonological structure, the relevant distinctions should be represented in overlapping brain regions for both pseudowords and real words. Such anatomical convergence would suggest that similar mechanisms contribute to iconic sound-meaning mappings across stimulus types. However, real words differ from pseudowords because their sound-based cues complement learned semantic representations. As a result, semantic knowledge may shape how acoustic and phonological cues are processed, leading to both shared and stimulus-specific patterns of neural processing. We therefore hypothesize that neural processes associated with iconicity partially co-localize across pseudowords and real words, consistent with a shared acoustic-phonetic basis that may be modulated by semantic knowledge in the case of real words.

To test this hypothesis, we designed an fMRI study in which participants listened to real words and pseudowords and judged whether each stimulus sounded rounded or pointed. The pseudowords were generated using phonemes from the real words and constructed to preserve their acoustic, phonetic and phonotactic properties, while lacking established lexical meanings. Using multivariate decoding, we tested whether rounded and pointed items could be distinguished by examining the distributed patterns of brain activity, separately for pseudowords and real words. We examined whether brain regions showing sensitivity to rounded-pointed distinctions for pseudowords and real words converged spatially. We further investigated whether classifiers trained to distinguish rounded from pointed real words could also distinguish rounded/pointed pseudowords, and vice versa, to assess whether discriminative information generalized across stimulus types. This approach allowed us to assess both shared and differential contributions to iconicity.

## 2. METHODS

### a. Participants

Thirty-two participants took part in the fMRI study. Four participants were excluded from the analyses: one discontinued the scan before completing the session, one had no recorded behavioral responses due to a response-box malfunction, one failed to meet the criterion for above-chance (see section 2d) behavioral performance and one had a perivascular-space abnormality identified on MRI. The reported results are therefore based on 28 participants (19 females; mean age = 29 years 2 months, SD = 6 years 2 months). All participants were right-handed according to the modified Edinburgh Handedness Inventory (Raczkowski et al., 1974). Participants reported normal hearing and normal or corrected-to-normal vision. All participants provided written informed consent, and the study was approved by the Penn State University Institutional Review Board.

### b. Stimuli

We recorded two sets of auditory items: real words and pseudowords. The real words were selected based on their roundedness or pointedness scores from the list of Sidhu et al. (2021), and were all bisyllabic. To generate pseudowords, we recombined the phonemes within each of the pointed and rounded real word sets while preserving the consonant-vowel structure of the original words, resulting in a set of “rounded” and another set of “pointed” pseudowords (**Figure 1A**). This ensured that the phonological characteristics associated with roundedness and pointedness were preserved across stimulus types. Using this procedure, we initially created 48 bisyllabic pseudowords, consisting of 24 rounded and 24 pointed candidates.

**Figure 1:**
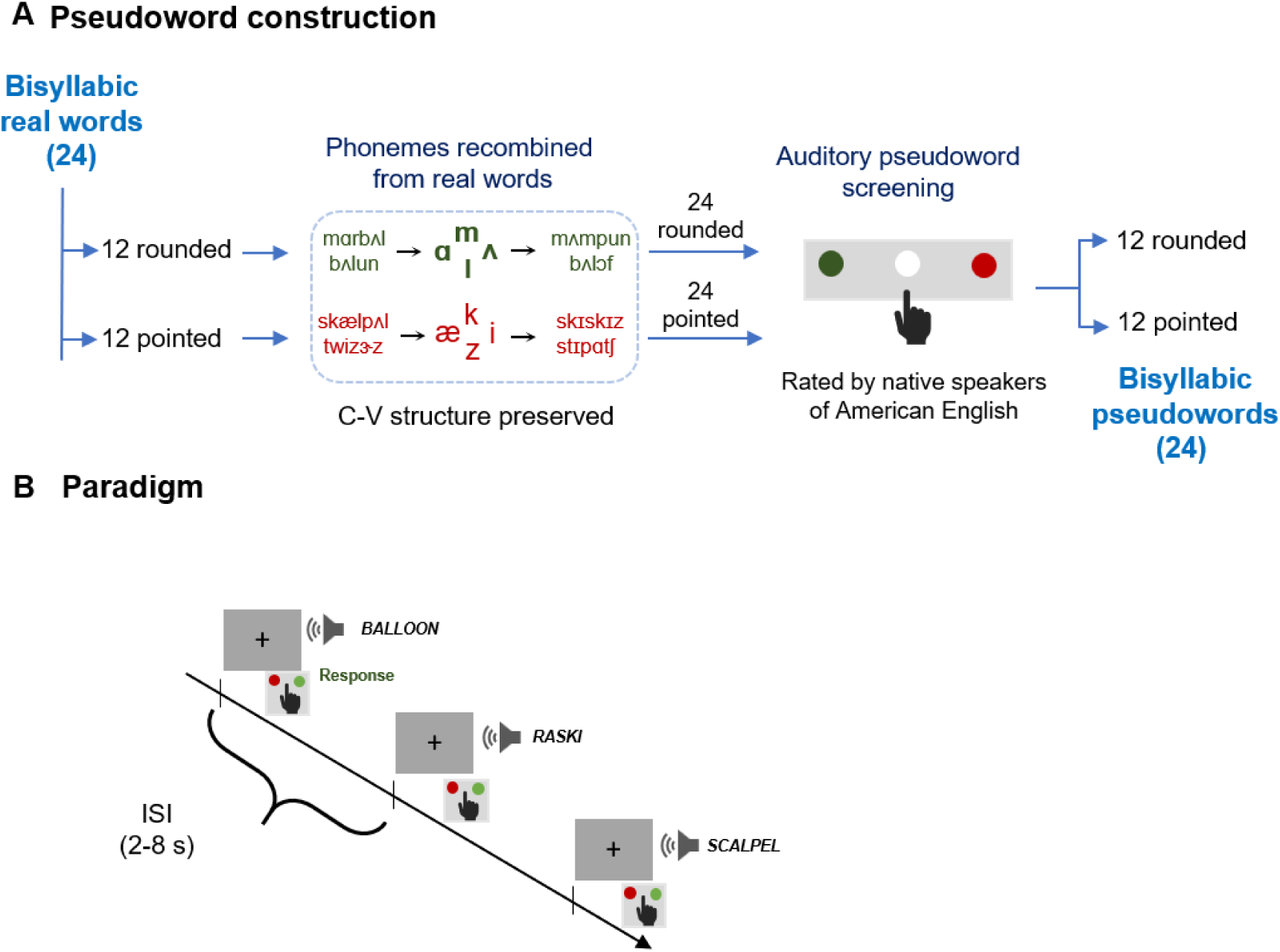
Stimulus construction and experimental paradigm. **A.** Phonemes from 24 bisyllabic real words were recombined separately within each of the two semantic categories (rounded and pointed) to generate candidate pseudowords while preserving the consonant-vowel structure. Within each semantic category, the real-word and pseudoword sets were matched in overall phoneme content and phonotactic probability. **B.** The experiment comprised an auditory classification task during fMRI using a jittered event-related design spanning 8 runs. Participants heard a single item and indicated whether it sounded rounded or pointed via button press.

The 24 selected real words and the 48 pseudowords generated through phoneme recombination were recorded by a female native speaker of American English, who produced all items with neutral intonation in randomized order. The recordings were digitized at 44.1 kHz using Praat speech analysis software (Boersma & Weenick, 2012). The audio files were subsequently downsampled to 22.05 kHz and amplitude-normalized using Audacity v3.2 (Audacity Team, 2012). Three native speakers of American English then classified the auditory pseudowords as sounding rounded, pointed, or neutral. Based on these ratings, we selected the 12 pseudowords most consistently identified as rounded and the 12 most consistently identified as pointed for the final stimulus set. Thus, the final stimulus sets consisted of 24 real words and 24 pseudowords, each set containing 12 rounded and 12 pointed items. Finally, we ensured that the rounded and pointed pseudoword sets were similar to the corresponding real word sets in phoneme content: phonotactic probability did not differ significantly between real words and pseudowords (*t_23_* = 0.83, *p* = 0.41) (**Figure 1A**). The mean stimulus duration was 0.964s for real words (SD = 0.18s) and 1.074s for pseudowords (SD = 0.17s).

### c. Experimental procedure

#### In-scanner task

Participants completed an auditory classification task during fMRI using a jittered, event-related design. The experiment comprised eight functional runs, each containing 48 trials equally divided among four conditions: rounded and pointed real words, and rounded and pointed pseudowords. On each trial, participants listened to an auditory item and judged whether it *sounded* rounded or pointed (**Figure 1B**). Participants were instructed to respond based on their first intuition. Stimulus onsets were jittered, with interstimulus intervals ranging from 2 to 8s. Each run began and ended with a 10s rest period, resulting in a total run duration of 332s. Responses were made using the right index and middle fingers on a handheld response box, with the button mapping for rounded and pointed responses counterbalanced across participants. Accuracy and reaction times were recorded, and stimulus presentation and response collection were controlled using Presentation software (Neurobehavioral Systems, http://www.neurobs.com).

### d. Behavioral analyses

Behavioral performance was assessed for each participant using one-sided exact binomial tests against a chance level of 50% (Siegel & Castellan, 1988). Correct responses and valid trials were pooled across runs, separately for real words and pseudowords. Trials with missing responses were excluded. Participants were required to demonstrate significantly above-chance performance in both conditions at an uncorrected significance threshold of α = .05 to be included in further analyses. For each participant and run, accuracy was calculated as the percentage of correct responses. Accuracy was calculated separately for rounded and pointed stimuli and then averaged within each lexical condition (real words and pseudowords). Participant-level accuracy was obtained by averaging across runs. For group analyses, real-word and pseudoword accuracy were compared using a two-sided paired-samples t test.

### e. Imaging

MRI data were acquired at the Center for Nuclear Magnetic Resonance Imaging at Penn State College of Medicine using a 3T Siemens Prisma Fit scanner equipped with a 64-channel head/neck receive coil. Participants lay supine in the scanner with foam padding placed around the head to minimize movement. Siemens three-dimensional prospective acquisition correction (3D PACE) was implemented to reduce motion-related artifacts during scanning. Auditory stimuli were presented through MRI-compatible high-fidelity headphones (Sensimetrics, Malden, MA), which also attenuated ambient scanner noise.

T1-weighted anatomical images were collected using a 3D MPRAGE sequence (repetition time (TR) = 2.30 s, echo time (TE) = 2.34 ms, inversion time (TI) = 807 ms, flip angle = 9°). Images were acquired with 1 mm isotropic voxel resolution, yielding a field of view (FOV) of 176 × 256 × 256 mm. Parallel imaging was implemented using GRAPPA with an acceleration factor of 2.

Functional images were acquired using a T2*-weighted gradient-echo echoplanar imaging (EPI) sequence. Whole-brain volumes were obtained with repetition time (TR) = 2.0 s, echo time (TE) = 30 ms, and flip angle = 90°. Thirty-six axial slices were acquired with 3 mm isotropic voxels, resulting in an FOV of 192 × 192 × 108 mm. Each run consisted of 166 volumes. In-plane parallel imaging was applied using GRAPPA (acceleration factor = 2). Phase encoding was performed in the posterior-to-anterior direction (j−). Once magnetic stabilization was achieved at the start of each run, the scanner triggered the computer running Presentation software, ensuring synchronization between experimental stimulus delivery and image acquisition.

### f. MRI data analyses

#### Data preprocessing

MRI data were preprocessed using fMRIPrep version 24.1.0 (Esteban et al., 2019, 2020), an automated pipeline built on Nipype (Gorgolewski et al., 2011). Functional and anatomical images were organized according to the Brain Imaging Data Structure (BIDS) prior to preprocessing and processed on a high-performance computing cluster. Structural preprocessing began with correction of spatial intensity inhomogeneities in the T1-weighted anatomical image using N4 bias-field correction (Tustison et al., 2010) implemented in the Advanced Normalization Tools (ANTs) library (Avants et al., 2008). The corrected T1-weighted image served as the anatomical reference volume for all subsequent preprocessing steps. Following intensity correction, the anatomical image was skull-stripped to remove non-brain tissue. The brain-extracted anatomical image was then segmented into gray matter (GM), white matter (WM), and cerebrospinal fluid (CSF) using tissue segmentation algorithms implemented within the preprocessing workflow. These tissue probability maps were used to generate subject-specific anatomical masks.

Functional MRI data consisted of eight BOLD runs per participant. For each run, preprocessing followed the standard fMRIPrep functional workflow. Functional images were corrected for slice-timing differences and head motion using rigid-body transformations that estimated six motion parameters describing translation and rotation of the head across volumes. The motion-corrected functional images were then co-registered to the participant’s anatomical image using boundary-based registration, ensuring accurate alignment between functional and structural data. Several nuisance regressors were estimated during preprocessing to characterize potential sources of non-neural variance in the BOLD signal. These included head-motion parameters, framewise displacement, and physiological signals derived from tissue masks. Global signals from white matter, cerebrospinal fluid, and the whole brain were extracted and stored alongside the preprocessed functional data. These confound time series were used in subsequent analyses to account for motion-related and physiological noise.

#### fMRI data analysis

Neural responses were estimated using general linear models (GLMs) implemented in Statistical Parametric Mapping software, version 12 (SPM12; Wellcome Trust Centre for Neuroimaging, London, UK). First-level model estimation was conducted separately for the univariate and multivariate analyses because the two analyses differed in when spatial smoothing was applied. For univariate analysis, the preprocessed BOLD time series was spatially smoothed in native space using an 8 mm full-width at half-maximum (FWHM) isotropic Gaussian kernel before first-level model estimation. For multivariate analyses, the preprocessed BOLD time series was not spatially smoothed prior to first-level model estimation, to preserve fine-grained spatial information for multivariate decoding. Instead, smoothing was applied later to the resulting searchlight decoding maps before group-level analysis as described in the multivariate pattern analysis section below.

For both analyses, first-level models were estimated using a trial-wise GLM, in which each stimulus presentation was modeled with a separate regressor and convolved with the canonical hemodynamic response function. Each functional run contained 48 trials, resulting in 48 trial-specific regressors per run. This approach provided stimulus-specific response estimates for both univariate and multivariate analyses. To account for motion-related variance, six rigid-body motion parameters derived from the fMRIPrep confounds were included as nuisance regressors. Temporal autocorrelation in the BOLD time series was modeled using the AR(1) autocorrelation model implemented in SPM12, and low-frequency drifts were removed using a high-pass filter with a cutoff period of 128 s. Model parameters were estimated using the classical ordinary least-squares approach implemented in SPM12. An explicit analysis mask derived from the fMRIPrep brain mask was applied to restrict estimation to within-brain voxels.

##### Univariate analysis

Univariate analyses were conducted to assess whether iconic associations were reflected in differences in the BOLD signal evoked by the stimuli. To examine whether activation differences associated with iconicity were present within each item type, separate linear contrasts were computed comparing the neural responses to rounded and pointed items, separately for real words and pseudowords. We also tested whether the magnitude of the iconicity effect differed between real words and pseudowords by computing an interaction contrast.

The resulting subject-level contrast maps were normalized to standard MNI space using nonlinear transformations estimated during preprocessing and implemented with ANTs (Avants et al., 2008), and then entered into group-level one-sample *t*-test analyses, treating subjects as a random factor. An explicit GM mask derived during preprocessing was applied at the group level to restrict analyses to GM voxels. Correction for multiple comparisons was performed using topological false discovery rate (FDR) correction, with clusters defined at an initial voxel-wise threshold of *p* < 0.001 and retained at a cluster-level FDR threshold of *q* < 0.05 (Chumbley et al., 2010). These contrasts allowed us to evaluate whether iconic associations were reflected in activation differences within real words and pseudowords, and whether such effects differed as a function of word type.

##### Multivariate pattern analysis (MVPA)

MVPA was performed using The Decoding Toolbox (Hebart et al., 2015) together with SPM12. The analysis used the unsmoothed trial-wise beta estimates obtained from the first-level GLM. These beta images were used directly for decoding to preserve fine-grained spatial information relevant for MVPA. A searchlight decoding approach was implemented to determine whether local spatial patterns of activity could discriminate rounded versus pointed items. For each voxel, a spherical neighborhood with a 4-voxel radius (12 mm) was defined, and the multivoxel activity pattern within that sphere was used for classification. Decoding analyses were conducted separately for real words and pseudowords. Cross-classification analyses were also conducted in both directions: training the classifier to distinguish rounded versus pointed pseudowords and testing it on real words, and training it on real words and testing it on pseudowords. Classification was performed using a support vector machine classifier implemented through the LIBSVM package (Chang & Lin, 2011). In order to ensure independence between training and testing data, decoding used a leave-one-run-out cross-validation scheme. In each iteration, data from seven runs were used to train the classifier, and data from the remaining run were used for testing. This procedure was repeated until each run served once as the independent test set. For each searchlight location, decoding performance was quantified as classification accuracy relative to chance level for each participant, separately for real words and pseudowords.

Searchlight analyses were restricted to voxels within a subject-specific brain mask derived from the first-level analysis. The MVPA yielded four native-space classification accuracy searchlight maps for each participant, corresponding to rounded versus pointed classification separately for real words and pseudowords and cross-classification in both directions (training on pseudowords and testing on real words, and training on real words and testing on pseudowords). Individual searchlight classification accuracy maps were smoothed with a 4 mm FWHM isotropic Gaussian kernel and normalized to MNI space using nonlinear transformations estimated during preprocessing and implemented with ANTs (Avants et al., 2008). Group-level one-sample t-tests were then conducted separately for real words and pseudowords, treating subjects as a random factor. An explicit GM mask was applied to restrict analyses to GM voxels. Statistical significance at the group level was assessed using topological false discovery rate correction, with clusters defined at an initial voxel-wise threshold of p < 0.001 and retained at a cluster-level FDR threshold of q < 0.05 (Chumbley et al., 2010).

#### Robustness analysis of MVPA

To ensure that the observed decoding effects were not dependent on specific analysis parameters or individual participants, robustness was assessed across both spatial scale and participant inclusion (Etzel et al., 2013). To assess robustness across spatial scale, the same searchlight decoding procedure was repeated using additional radius lengths of 6 mm and 9 mm. For each radius, group-level statistical analysis was performed using topological FDR correction as outlined above. Robustness across spatial scale was evaluated by examining whether the main clusters identified in the 12 mm-radius analysis were preserved in the analyses with the smaller radii. Participant-level robustness was assessed by repeating the same group-level analysis iteratively while leaving out one participant at a time. The resulting statistical maps were manually inspected to determine whether the significant clusters observed in the primary 12 mm analysis were consistently present across leave-one-participant-out iterations. These procedures confirmed that the reported decoding effects were not driven by a single participant or by the specific choice of searchlight radius.

## 3. RESULTS

### a. Behavior

Following the exclusions described in the Methods, 28 participants were included in the analyses. Mean classification accuracy was 88.7% (SD = 8.9%) for real words and 82.6% (SD = 9.9%) for pseudowords (**Figure 2**). A paired *t-*test comparing accuracy for real words and pseudowords across participants showed that accuracy was significantly higher for real words compared to pseudowords (*t_27_* =3.11, *p* = 0.004). All participants performed significantly above chance in both conditions (**Supplementary Figure 1**).

**Figure 2:**
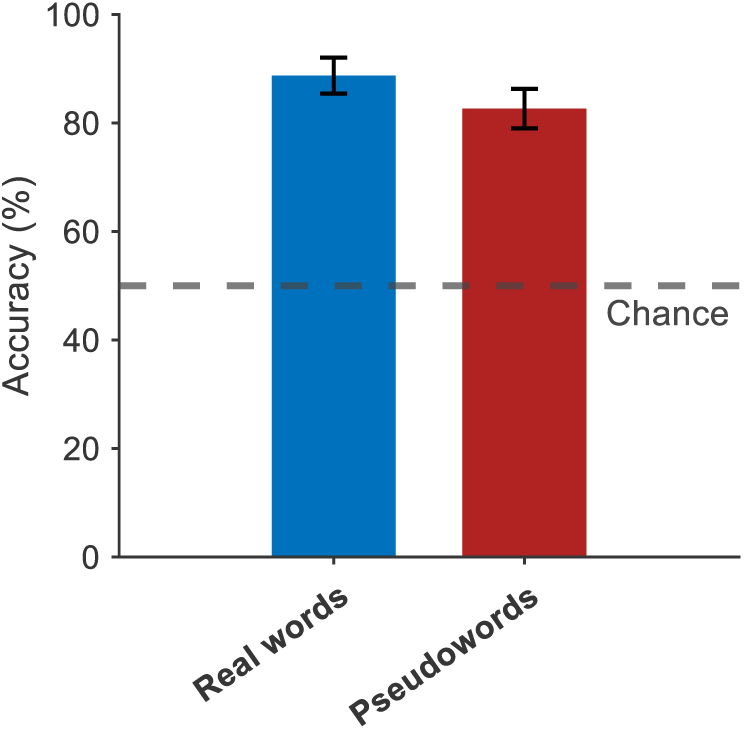
Behavioral results. Participants performed a two-alternative forced-choice task during fMRI, classifying auditory stimuli as “rounded” or “pointed”. Colored bars represent mean classification accuracy across participants (N = 28), with error bars indicating ± SEM. The dashed line indicates chance-level performance (50%).

### b. Univariate analyses

After correction for multiple comparisons, no significant clusters were observed for the rounded versus pointed contrast either for real words or pseudowords, nor for the interaction contrast testing whether the magnitude of differences for rounded versus pointed items differed between real words and pseudowords. The absence of significant univariate effects indicates that iconic associations between rounded and pointed items were not reflected in the mean differences in BOLD activation. Thus, BOLD responses to iconic associations were not expressed as simple increases or decreases in activity of voxels, motivating the use of multivariate analyses to test whether these distinctions are instead reflected in distributed spatial patterns of activity.

### c. MVPA

Searchlight decoding was employed to examine whether rounded versus pointed iconic associations could be discriminated in local patterns of BOLD activity within real words and pseudowords. Cross-classification tested whether this distinction generalized across lexical conditions by training classifiers on real words and testing on pseudowords, and vice versa. Group-level analyses revealed significant above-chance decoding in all four analyses, indicating that patterned information distinguishing rounded from pointed items was present for both real words and pseudowords (**Figure 3**, **Table 1**), and that this distinction generalized across lexical conditions in both directions within specific cortical regions (**Figure 4**, **Table 2**).

**Figure 3:**
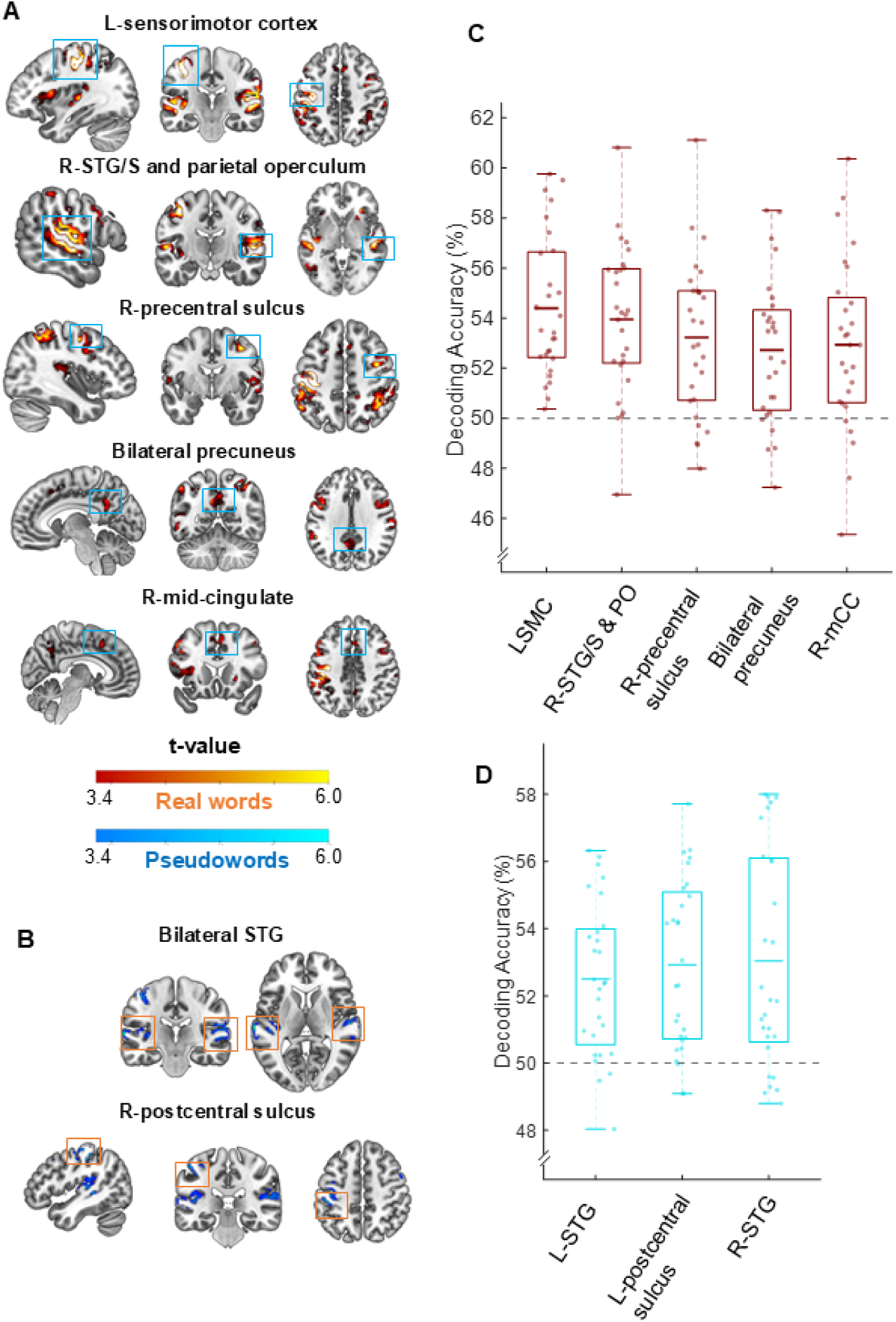
Within-condition searchlight decoding of rounded versus pointed real words and pseudowords. Searchlight decoding results for (A) real words (orange) and (B) pseudowords (cyan). Statistical maps show clusters surviving correction for multiple comparisons. Box plots show mean decoding accuracy (%) extracted from the peak voxel of each significant cluster for (C) real words and (D) pseudowords. In the box plots, the horizontal line within the box indicates the mean, the box represents the interquartile range, and the whiskers show the spread of values outside the interquartile range. Individual data points represent participants. L: left hemisphere; R: right hemisphere STG: Superior temporal gyrus; STS: Superior temporal sulcus; PO: parietal operculum; SMC: Sensorimotor cortex; mCC: mid-cingulate cortex

**Figure 4:**
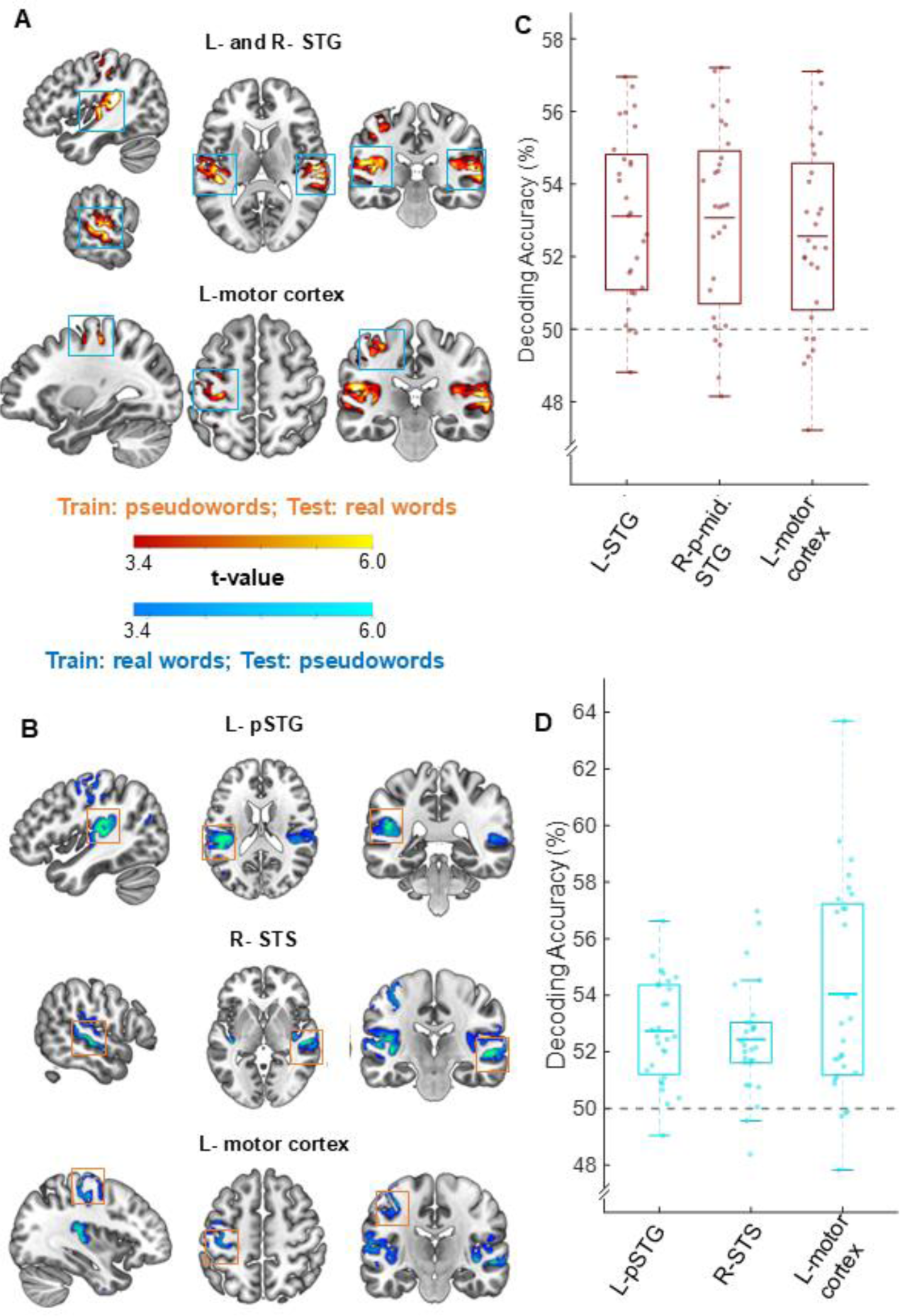
Cross-condition searchlight decoding of rounded versus pointed items. Searchlight maps show significant clusters for classifiers: (A) trained on pseudowords and tested on real words (orange), and (B) trained on pseudowords and tested on real words(cyan). Box plots show participant-level decoding accuracy (%) extracted from the peak voxel of each significant cluster for (C) pseudoword-to-real-word direction and (D) real-word-to-pseudoword direction. Details as in Fig. 3. L: left hemisphere; R: right hemisphere STG: Superior temporal gyrus; STS: Superior temporal sulcus

**Table 1:**
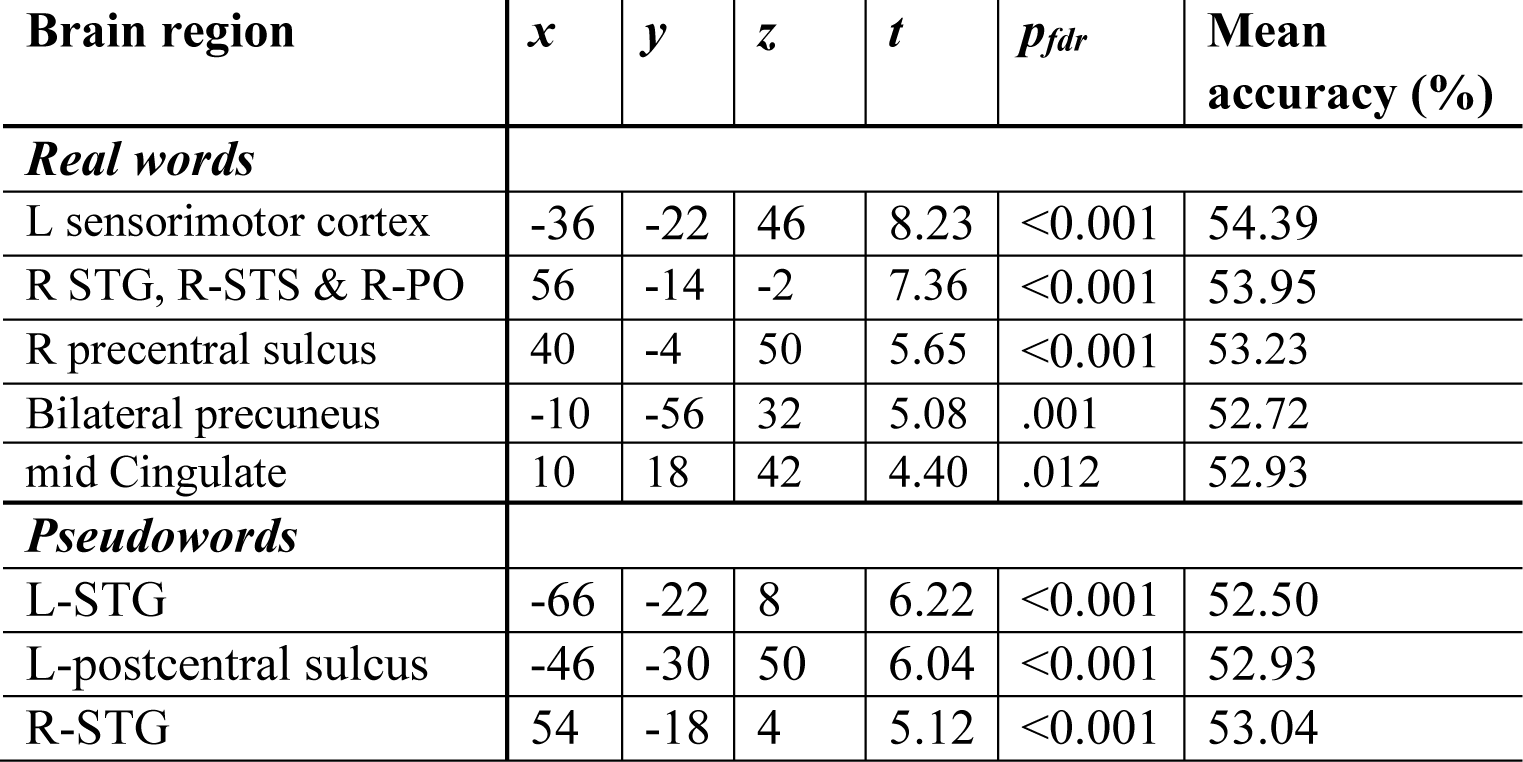
Significant clusters from within-condition searchlight decoding of iconic associations. Coordinates (x, y, z; MNI space) indicate the peak voxel of each cluster. Reported t-values, corrected p-values, and mean decoding accuracy (%) were extracted at the peak voxel. Statistical significance was assessed using an initial voxel-wise cluster-forming threshold of p < .001, followed by cluster-level topological false discovery rate correction at q < .05. L: left hemisphere; R: right hemisphere STG: Superior temporal gyrus; STS: Superior temporal sulcus; PO: parietal operculum

| **Brain region** | ***x*** | ***y*** | ***z*** | ***t*** | ***p<sub>fdr</sub>*** | **Mean accuracy (%)** |
|---|---|---|---|---|---|---|
| ***Real words*** |  |  |  |  |  |  |
| L sensorimotor cortex | -36 | -22 | 46 | 8.23 | <0.001 | 54.39 |
| R STG, R-STS & R-PO | 56 | -14 | -2 | 7.36 | <0.001 | 53.95 |
| R precentral sulcus | 40 | -4 | 50 | 5.65 | <0.001 | 53.23 |
| Bilateral precuneus | -10 | -56 | 32 | 5.08 | .001 | 52.72 |
| mid Cingulate | 10 | 18 | 42 | 4.40 | .012 | 52.93 |
| ***Pseudowords*** |  |  |  |  |  |  |
| L-STG | -66 | -22 | 8 | 6.22 | <0.001 | 52.50 |
| L-postcentral sulcus | -46 | -30 | 50 | 6.04 | <0.001 | 52.93 |
| R-STG | 54 | -18 | 4 | 5.12 | <0.001 | 53.04 |

**Table 2:**
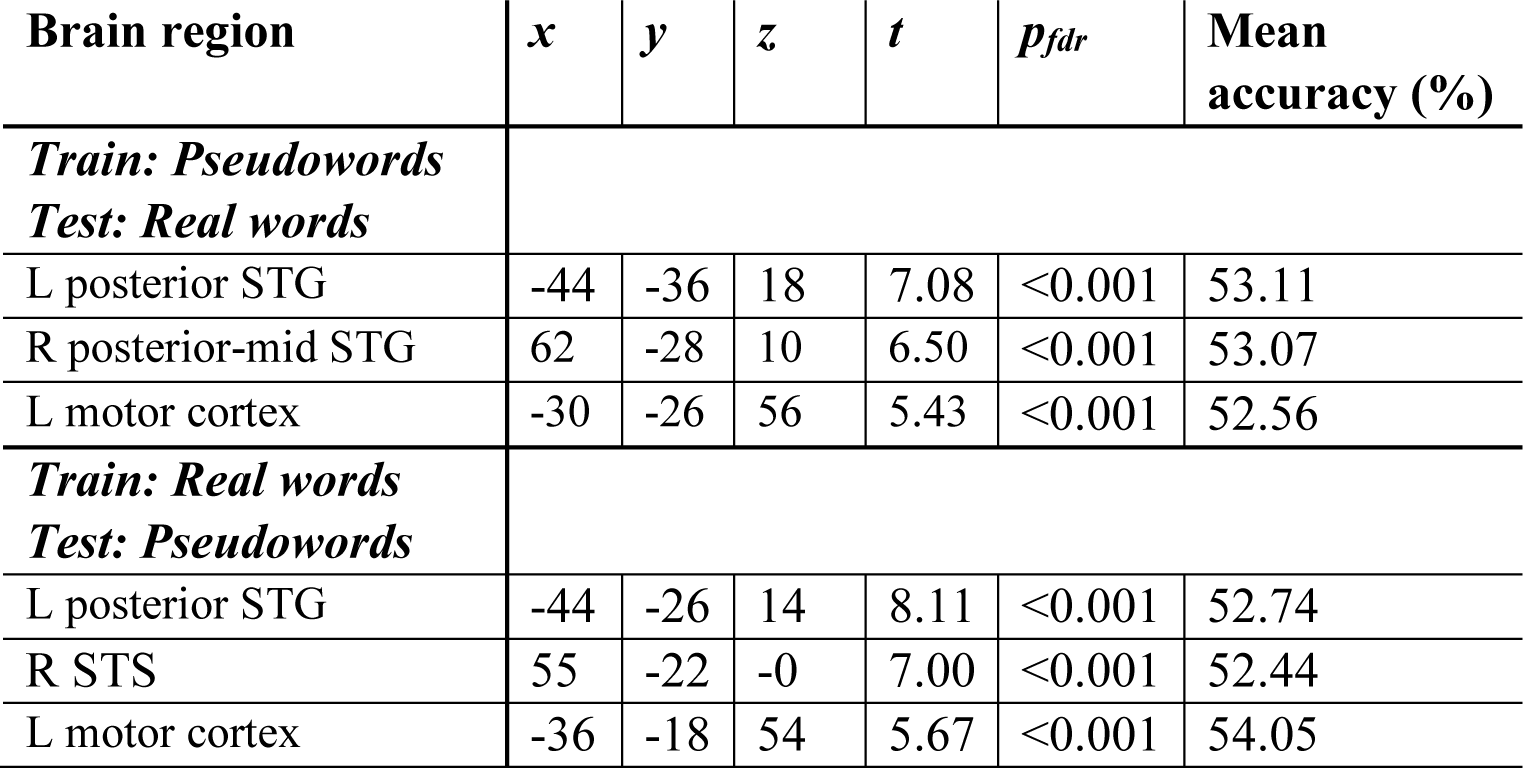
Significant clusters from cross-condition searchlight decoding of iconic associations. . Details as in Table 1.

| **Brain region** | ***x*** | ***y*** | ***z*** | ***t*** | ***p<sub>fdr</sub>*** | **Mean accuracy (%)** |
|---|---|---|---|---|---|---|
| ***Train: Pseudowords***<br>***Test: Real words*** |  |  |  |  |  |  |
| L posterior STG | -44 | -36 | 18 | 7.08 | <0.001 | 53.11 |
| R posterior-mid STG | 62 | -28 | 10 | 6.50 | <0.001 | 53.07 |
| L motor cortex | -30 | -26 | 56 | 5.43 | <0.001 | 52.56 |
| ***Train: Real words***<br>***Test: Pseudowords*** |  |  |  |  |  |  |
| L posterior STG | -44 | -26 | 14 | 8.11 | <0.001 | 52.74 |
| R STS | 55 | -22 | -0 | 7.00 | <0.001 | 52.44 |
| L motor cortex | -36 | -18 | 54 | 5.67 | <0.001 | 54.05 |

#### Within-condition decoding

For real words, significant decoding of rounded versus pointed items was observed in a distributed set of cortical regions (**Figure 3A, C**). The strongest effect was observed in the left sensorimotor cortex (L-SMC; *t* = 8.23, *p_fdr_* < .001; mean accuracy = 54.4%), followed by the right superior temporal gyrus/sulcus extending into the parietal operculum (R-STG/STS–PO; *t* = 7.36, *p_fdr_* < .001; mean accuracy = 53.9%). Additional significant clusters were identified in the right precentral sulcus (*t* = 5.65, *p_fdr_* < .001; mean accuracy = 53.2%), bilateral precuneus (*t* = 5.08, *p_fdr_* = 0.001; mean accuracy = 52.7%) and right mid-cingulate cortex (R-mCC; *t* = 4.40, *p_fdr_* = 0.012; mean accuracy = 52.9%). Robustness analyses indicated that most real-word decoding effects were preserved across searchlight radii and participant inclusion. Of the clusters identified in the primary 12 mm-radius analysis, the L-SMC, R-STG/STS–PO, and R-mCC remained significant in both the 9 mm- and 6 mm-radius analyses. The right precentral sulcus cluster remained significant at 6 mm but did not survive correction at 9 mm, whereas the bilateral precuneus cluster remained significant at 9 mm but did not survive correction at 6 mm. In leave-one-participant-out analyses, all clusters except the R-mCC were retained in at least 96% of iterations. The R-mCC cluster was retained in 57% of iterations, indicating lower stability across participant subsets despite its consistency across searchlight radii.

For pseudowords, significant decoding of rounded versus pointed items was observed in three clusters (**Figure 3B, D**): the left superior temporal gyrus (L-STG; *t* = 6.22, *p_fdr_* < .001; mean accuracy 52.5%); left postcentral sulcus (*t* = 6.04, *p_fdr_* < .001; mean accuracy = 52.9%) and right superior temporal gyrus (R-STG; *t* = 5.12, *p_fdr_* < .001; mean accuracy = 53%). Robustness analyses indicated that pseudoword decoding effects were largely preserved across searchlight radii and participant inclusion. Both STG clusters remained significant in the 9 mm- and 6 mm-radius analyses. The left postcentral sulcus cluster remained significant at 9 mm but did not survive correction at 6 mm. In leave-one-participant-out analyses, all three clusters were retained in every iteration, indicating that the effects were not driven by any single participant.

#### Cross-condition decoding

Cross-condition decoding revealed significant generalization of rounded versus pointed distinctions in both directions (**Figure 4**, **Table 2**). When classifiers were trained on pseudowords and tested on real words (**Figure 4A, C**), significant decoding was observed in the left posterior superior temporal gyrus (L-pSTG; *t* = 7.08, *p_fdr_* < .001; mean accuracy = 53.1%), right posterior-to-middle superior temporal gyrus (R-pSTG; *t* = 6.50, *p_fdr_* < .001; mean accuracy = 53.1%) and left motor cortex (*t* = 5.43, *p_fdr_* < .001; mean accuracy = 52.6%).

When classifiers were trained on real words and tested on pseudowords (**Figure 4B, D**), significant decoding was observed in the L-pSTG (L-pSTG; *t* = 8.11, *p_fdr_* < .001; mean accuracy = 52.7%), right superior temporal sulcus (R-STS; *t* = 7.00, *p_fdr_* < .001; mean accuracy = 52.4%) and left motor cortex (*t* = 5.67, *p_fdr_* < .001; mean accuracy = 54%). Thus, cross-condition decoding was observed in bilateral superior temporal regions and left motor cortex in both directions, indicating that local activity patterns carried information distinguishing rounded from pointed items that generalized across lexical conditions.

Robustness analysis indicated that cross-condition decoding was highly stable in both directions. All clusters identified in the primary 12 mm-radius analysis remained significant in both the 9 mm- and 6 mm-radius analyses. In leave-one-participant-out analyses, all clusters were retained in more than 95% of iterations in each direction, demonstrating consistency across searchlight radii and participant subsets.

## 4. DISCUSSION

Iconicity provides an important window into how sound and meaning are linked in natural language. Accumulating evidence highlights the ubiquity of iconicity across languages (Blasi et al., 2016) and its functional significance in language learning (Imai & Kita, 2014; Kantartzis et al., 2019) and development (Maurer et al., 2006; Ozturk et al., 2013). Therefore, studying iconicity in spoken words is critical for understanding how sound cues are mapped onto meaning during speech perception. However, much of what we know about iconic sound-meaning associations comes from studies using engineered pseudowords, which have been valuable for identifying the acoustic parameters (Knoeferle et al., 2017; Kumar et al., 2025; Lacey et al., 2026), linguistic features (Cuskley et al., 2017; McCormick et al., 2015; Nielsen & Rendall, 2013), and neural processes (Barany et al., 2023; McCormick et al., 2022; Peiffer-Smadja & Cohen, 2019) that mediate these associations. Less is known about whether the same neural mechanisms operate when sound cues are embedded within real words, which carry established semantic knowledge in addition to iconic cues. Therefore, by comparing real words and pseudowords matched in acoustic, phonemic, and phonotactic properties, the present study examined whether the neural mechanisms that mediate sound-to-meaning mapping in pseudowords also extend to natural language and whether neural processing differs in the presence or absence of semantic knowledge. In addition to decoding rounded versus pointed items separately for each stimulus type, we used bi-directional cross-classification to test whether neural patterns distinguishing these categories generalize across stimulus type.

We reasoned that if iconic mappings are strongly shaped by acoustic and phonetic structure, then real words and pseudowords should exhibit spatial convergence in the cortical loci that show significant decoding accuracy for rounded versus pointed items. Such convergence would suggest that similar mechanisms contribute to iconic sound-meaning mappings across stimulus types. To assess whether neural information was shared across stimulus types, we used cross-classification to test whether neural patterns distinguishing rounded from pointed items generalized between real words and pseudowords.

We first confirmed that participants showed robust sensitivity to rounded/pointed iconic associations for both real words and pseudowords. Accuracy was high for both stimulus types (**Figure 2**), indicating that participants were able to use the auditory properties of the stimuli to make reliable rounded versus pointed judgments. This confirms that the stimulus sets captured iconic associations. Importantly, we observed that behavioral classification accuracy was significantly higher for real words than for pseudowords. This suggests that although acoustic and phonetic cues were more than sufficient to support iconic judgments for pseudowords, the presence of established meaning in real words may have strengthened these judgments. Importantly, the higher accuracy for real words suggests that when a word’s sound and meaning evoke the same shape category, these cues may reinforce one another and support more accurate rounded versus pointed judgments.

### Mean BOLD responses

The univariate analyses showed no significant activation differences for rounded versus pointed items for either real words or pseudowords. The interaction contrast also did not reveal evidence that the magnitude of any rounded-pointed activation differences varied between stimulus types. This contrasts with previous fMRI studies using explicit and implicit sound-shape association tasks, where univariate effects were linked to congruency, multisensory attention or conflict processing (Barany et al., 2023; McCormick et al., 2022; Peiffer-Smadja & Cohen, 2019). In the present study, participants heard only auditory stimuli and judged whether each item sounded rounded or pointed, allowing us to examine iconic associations for spoken items rather than responses to crossmodal matching. The lack of univariate effects therefore suggests that rounded versus pointed iconic distinctions are not primarily expressed as changes in BOLD activation, providing a basis for examining whether these distinctions are instead reflected in patterned BOLD responses, which can be detected using multivariate decoding.

### Patterned BOLD responses

MVPA showed that rounded and pointed items could be discriminated in local patterns of BOLD activity for both real words and pseudowords. This shows that iconic distinctions were present at the level of patterned responses.

### Real word decoding

For real words, significant decoding was observed across a distributed set of cortical regions (**Figure 3A, C**). Mean decoding accuracy at the cluster peaks ranged from 52.7% to 54.4% (**Table 1**). These values indicate statistically significant but modest discrimination between rounded and pointed items. Decoding in the right STG and STS, within a cluster extending into the parietal operculum, may reflect the processing of acoustic and phonetic properties associated with rounded and pointed judgments. Previous work has demonstrated that superior temporal activity patterns distinguish phonetic features of speech (Arsenault & Buchsbaum, 2015), including in the right hemisphere. More specifically, manipulating the spectral and temporal detail of spoken words revealed relatively greater sensitivity to spectral detail in right than left STS (Obleser et al., 2008). This provides a possible basis for considering the contribution of spectral properties to the right temporal cortex decoding observed here. The extension into the parietal operculum may reflect sensitivity to sound frequency, given evidence that somatosensory activity patterns distinguish tones of different frequencies (Pérez-Bellido et al., 2018), although the contiguity with adjacent auditory cortex leaves this uncertain. Therefore, our findings are consistent with speech-signal differences contributing to the discrimination of real word categories, although decoding alone does not identify which acoustic properties contributed.

The highest peak decoding accuracy occurred in the left sensorimotor cortex, while significant decoding was also observed in the right precentral sulcus. Individual finger movements elicit distinguishable activity patterns in primary motor and somatosensory cortex (Ariani et al., 2022; Yokoi et al., 2018), suggesting that the manual responses used to report the judgments may have contributed to these findings. Participants indicated their judgments using the index and middle fingers, with the response mapping counterbalanced across participants. This counterbalancing does not eliminate response-related information from within-participant decoding when each category retains a consistent finger mapping. The observed discrimination may therefore reflect the responses associated with rounded and pointed judgments. Consistent with this interpretation, activity in the precentral sulcus could reflect sensorimotor processes involved in the preparation of a response. Whether these regions also contribute to forming the iconic associations themselves cannot be determined from the present results.

Significant decoding bilaterally in the precuneus may reflect visual imagery during real-word judgments. Ganis et al. (2004) reported bilateral precuneus engagement when participants visualized previously studied drawings cued by spoken object names and judged their visual properties, including whether they contained circular or rectangular parts. Participants in the present study may similarly have generated mental images of rounded or pointed shapes when evaluating the words.

Significant decoding was also observed in the R-mCC. Functional connectivity analyses have implicated its anterior subdivision in cognitive motor control (Hoffstaedter et al., 2014). Consistent with this account, Hoffstaedter et al. (2013) found that anterior mid-cingulate activity increased when participants selected both which hand to move and when to initiate the movement. To the extent that the present cluster involves this subdivision, decoding may reflect processes involved in selecting and initiating the responses used to report rounded and pointed judgments.

### Pseudoword decoding

For pseudowords, rounded and pointed items could be distinguished from activity patterns in bilateral STG. STG is sensitive to the spectral and temporal structure of speech, with neural responses tracking acoustic cues associated with phonetic features (Hullett et al., 2016; Mesgarani et al., 2014). Superior temporal activity patterns also carry information about voicing, manner and place of articulation (Arsenault & Buchsbaum, 2015). Our bilateral STG findings may reflect overlapping but partly different contributions of the two hemispheres to acoustic–phonetic processing. Left STG decoding is consistent with sensitivity to acoustic cues that distinguish phonetic features (Mesgarani et al., 2014). In a direct comparison of hemispheres, Arsenault and Buchsbaum (2015) found moderately stronger left-hemisphere pattern information for place of articulation, although voicing and manner information were bilaterally represented. Right STG decoding could also reflect phonetic distinctions, with a possible additional contribution from relative sensitivity to spectral structure. Evidence from manipulated speech supports relatively greater right temporal sensitivity to spectral detail and greater left temporal sensitivity to temporal detail (Flinker et al., 2019; Obleser et al., 2008). These findings suggest possible differences in how the two hemispheres contribute to the judgments. Auditory pseudoword identity has also been decoded in bilateral STG and STS across changes in the accompanying visual shape (Barany et al., 2023). Together, these findings suggest that decoding in the STG could be associated with acoustic and phonetic differences between rounded and pointed pseudowords that contribute to listeners’ shape judgments. Significant decoding was also observed in the left postcentral sulcus, which may relate to the manual responses used to report shape judgments. Participants used different fingers to indicate rounded and pointed judgments, and previous work has demonstrated finger-specific activity patterns in primary somatosensory cortex during both movement preparation and execution (Ariani et al., 2022). Therefore, decoding in this region could reflect differences between the finger responses associated with the two judgments. Another possibility is that the left postcentral sulcus may be involved crossmodally in temporal judgments; we cannot currently rule out either possibility.

### Generalization of rounded and pointed distinction across real words and pseudowords

Cross-decoding revealed that classifiers trained to distinguish rounded from pointed pseudowords could also distinguish these categories in real words, and vice versa. Both directions yielded significant decoding in left posterior STG, right STG/S and left motor cortex. These findings extend the separate decoding analyses by showing that that some of the neural information distinguishing rounded from pointed pseudowords also distinguishes rounded from pointed real words. Generalization in STG/S is consistent with its sensitivity to acoustic and phonetic features in the spoken word (Arsenault & Buchsbaum, 2015; Hullett et al., 2016; Mesgarani et al., 2014). Because the pseudowords were constructed using real-word phonemes while maintaining phonotactic properties, successful transfer could reflect sensitivity to sound features associated with rounded and pointed judgments across both stimulus sets. This provides evidence that some information supporting classification is shared across real words and pseudowords. Generalization could reflect acoustic and phonetic similarities between items associated with the same shape category across the two stimulus sets. The present results therefore cannot determine whether the shared information is related to just the sounds of the items, their associated shapes, or both. Cross-decoding was also observed in left motor cortex. Activity patterns in this region can distinguish individual finger movements (Yokoi et al., 2018). The use of the same finger mapping for rounded and pointed judgments across real words and pseudowords for each participant could contribute to generalization between the two stimulus types. Therefore, significant cross-decoding in motor cortex could reflect the consistency of responses used to report these judgments.

### Shared information across real words and pseudowords

An important aim of this study was to determine whether real words and pseudowords showed similar cortical patterns for rounded versus pointed classification. Although both stimulus types showed reliable decoding, the distributions of significant decoding were not identical. While acoustic and phonetic information could contribute to iconic shape judgments for both stimulus types, the mechanism through which this information informs the judgments may differ. For pseudowords, judgments could rely primarily on differences in acoustic properties that suggest roundedness or pointedness. For real words, these acoustic cues occur alongside learned meanings that may evoke visual imagery of their referents, providing an additional basis for evaluating shape. Therefore, iconic judgments of real words could draw from both the shape suggested by the acoustic properties of the word and the shape associated with its meaning. Together, these findings suggest that sound structure contributes to iconic judgments across both stimulus types, while meaning and imagery could provide additional information during real-word judgments.

Our observations have implications for models of speech perception. Iconic cues embedded in the acoustic and phonetic structure of speech may be sufficient to support associations with sensory properties such as roundedness and pointedness, without requiring an established lexical meaning. In real words, these sound-based associations may operate alongside learned semantic information, which could reinforce or modify the association suggested by the sound. Iconicity may therefore function as an additional cue during speech perception, operating alongside phonological, lexical, semantic, and contextual information.

## Acknowledgements

This work was supported by institutional funds provided to KS by Penn State College of Medicine. We acknowledge the Penn State College of Medicine MRI Core for its support with fMRI data acquisition.

## Author contributions

G. Vinodh Kumar contributed to conceptualization, methodology, formal analysis and writing the original draft. Simon Lacey contributed to conceptualization, project administration, and manuscript review and editing. Lynne Nygaard contributed to conceptualization and manuscript review and editing. K. Sathian contributed to conceptualization, methodology, supervision, funding acquisition, and manuscript review and editing.

## Data and code availability

The authors will make the data and analysis code publicly available after publication.

## Declaration of the use of AI

The authors used Claude (Anthropic) and ChatGPT (OpenAI) to check grammar and rephrase certain parts of Introduction and Discussion. They also used Claude to help troubleshoot technical errors encountered while running the analysis on a high-performance computing cluster.

