## Supplementary Material for "DECODING SHAPE ICONICITY OF WORDS AND PSEUDOWORDS: AN fMRI STUDY"

Corresponding author*:

K. Sathian
Department of Neurology
Penn State Health Milton S. Hershey Medical Center
Hershey, PA 17033-0859, USA


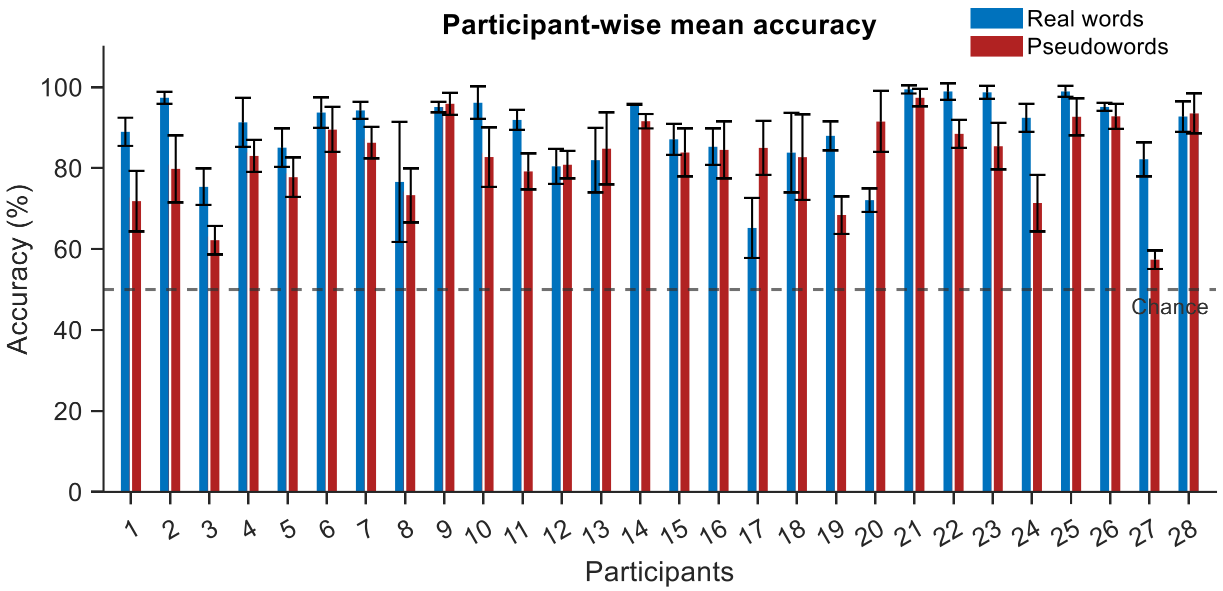


**Supplementary Figure 1. Behavioral performance for individual participants.** Mean accuracy for real words and pseudowords. The dashed line indicates chance-level performance (50%).
